# The Neural Impact Score benchmarks drugs in Multi-Region Brain Organoids

**DOI:** 10.64898/2026.08.26.746777

**Authors:** Aishwarya Pantula, Vanshita Singh, Om Sadul, Nitya Lagadapati, Kaustubh Joshi, Ryan Palaganas, Jon Sundstrom, Genevieve Stein-O’Brien, Annie Kathuria

## Abstract

Only about 10% of drugs that clear animal testing succeeds in humans, and central nervous system (CNS) programs carry an even steeper translational gap. Human-relevant NAMs are gaining global regulatory and funding support, creating an urgent need for interpretable preclinical systems that can generate comparable, decision-ready evidence across assays, models, and species. Yet the multimodal treatment-response data such systems produce are still evaluated assay by assay, with no unified metric showing whether a compound moves neural tissue toward a desirable or undesirable state. Here we present a framework called the Neural Impact Score (NIS), which translates multimodal CNS drug-response data into a bidirectional score across four pre-defined biological categories: neurodevelopment, neuroinflammation, neurodegeneration, and longevity. A positive score indicates a desirable shift, and a negative score indicates the opposite, placing compounds on a single scale across assays, model systems, and species. To demonstrate NIS, we analyzed a vascularized human day-200 multi-region brain organoid (MRBO) composed of cortical, endothelial, and brainstem lineages and mimicking a mid-gestational cortical window (GW18–GW22). We tested five compounds with distinct mechanisms of action: glucagon-like peptide-1 receptor agonist (GLP-1RA); norepinephrine-dopamine reuptake inhibitor (NDRI); selective serotonin reuptake inhibitor (SSRI); sphingosine-1-phosphate receptor modulator; and Akt activator, using single-nucleus RNA sequencing, bulk RNA sequencing, proteomics, and multi-electrode array electrophysiology. NIS integrated these readouts into category and composite scores, separating beneficial from adverse effects for each compound and sorting them into interpretation tiers. Applying the same framework to independent human and rodent datasets without retraining, we recovered conserved human antidepressant responses despite near-chance gene-level agreement between human MRBO and rat brain and identified an endothelial-dependent human-specific GLP-1 response absent from murine dorsal vagal complex. NIS therefore provides a human-relevant framework for drug evaluation and cross-species benchmarking, with a design extensible to other neural systems.

## Introduction

Global regulatory and funding agencies are increasingly prioritizing human-relevant New Approach Methodologies (NAMs)^1^ to improve the predictive relevance of nonclinical testing and reduce reliance on animal models. In the United States, initiatives from the US Food and Drug Administration (FDA) and National Institutes of Health ^2,3^ have encouraged human-based experimental systems, computational models, and other NAMs as part of a broader transition toward more predictive preclinical evidence. In Europe, EMA and the European Commission^4^ efforts similarly support regulatory acceptance of NAMs within the 3Rs framework, while the OECD’s international guidance emphasizes integrated approaches that combine in vitro, tissue-based, omics, and in silico evidence. This shift is also extending across Asia and Australia: Japan’s PMDA^5,6^ has identified NAMs as tools to improve prediction of human safety, efficacy, and pharmacokinetics; Korea’s MFDS^7^ and KoCVAM^8^ have advanced validated alternative methods into OECD Test Guidelines; China’s NMPA has promoted the development and adoption of alternative animal-testing methods in safety assessment; and Australia’s NHMRC^10^ and CSIRO^11^ have supported 3Rs implementation and national non-animal model capabilities for medical product development. Together, these initiatives are shifting preclinical research toward human-relevant systems that can generate evidence suitable for comparison, interpretation, and eventual decision-making.

This shift is especially urgent in CNS drug development, where animal studies poorly predict human outcomes and clinical attrition remains high. More than 90% of drugs^12^ that clear animal testing fails in humans, and CNS programs carry an even larger translational gap, including a 99.6% failure rate of Alzheimer’s disease candidates^13^ across clinical development. Human brain organoids and related NAM platforms can capture aspects of human neural biology that animal models miss, but biological complexity alone does not solve the translational problem. To support drug assessment, human-relevant models must generate interpretable, comparable, and decision-relevant outputs that can be evaluated across assays, compounds, model systems, and species.^14^

Single-region brain organoids have advanced human CNS modeling^15–18^, but they remain limited for drug-response assessment. Many lack vascular and blood-brain-barrier-associated compartments, do not capture interactions among multiple brain regions, and provide limited access to neurovascular or multi-compartment mechanisms of drug action. These limitations matter because CNS drug responses can involve molecular, cellular, vascular, and functional changes that do not localize to a single region or readout. Multi-region vascularized organoid platforms therefore offer an opportunity to improve human relevance by combining regional neural complexity with vascular context and scalable drug-response profiling.

We previously developed a vascularized human induced pluripotent stem cell (iPSC) derived Multi-Region Brain Organoid (MRBO) that integrated cortical, endothelial vascular, and brainstem lineages and aligns with a mid-gestational human cortical developmental window.^19^ This architecture provides a human tissue substrate for evaluating CNS drug responses across neural, glial, regional, and endothelial compartments rather than within a cortical-only model. However, generating multi-compartment biological data is only one part of the problem; these readouts also require a framework that can convert multimodal measurements into interpretable drug-response scores.

Multimodal profiling can measure CNS drug responses across transcriptomic,^20^ proteomic,^21^ and electrophysiological assays,^22–24^ but these readouts are often interpreted separately, leaving researchers to infer whether a compound shifts neural tissue toward favorable or unfavorable states. Previously developed factor models and weighted-nearest-neighbor approaches can identify latent components or joint embeddings,^25,26^ and connectivity-mapping tools can rank molecular similarity, but these outputs typically do not return signed biological-category scores that can be compared across assays, compounds, model systems, and species. This gap is especially important when human organoid responses must be compared with animal brain datasets that rarely contain matched assay modalities.

We developed the Neural Impact Score (NIS) to convert complex CNS drug-response data into a shared quantitative interpretation. NIS scores whether a compound supports neurodevelopment and plasticity, increases or suppresses neuroinflammation, protects against or promotes neurodegeneration, and increases or reduces cellular resilience. NIS integrates multi-omic readouts into bidirectional scores within predefined biological categories, where positive and negative values indicate favorable and unfavorable biological shifts, respectively. This allows compounds to be compared on the same scale across assay types and model systems. When full multimodal datasets are available, NIS generates integrated category-level and composite scores; when external comparator datasets are modality-restricted, the same locked category definitions can support cross-dataset and cross-species comparison.

In this study, we benchmarked NIS using healthy control human iPSC-derived MRBOs that we aged to 200 days for CNS drug assessment. Before disease-specific testing, healthy control models are needed to define baseline safety, tolerability, and early efficacy responses,^27^ and human MRBOs provide this first baseline in a developmental neural context that animal controls cannot fully capture.^28–30^ We profiled five CNS-relevant compounds spanning distinct mechanisms: a GLP-1 receptor agonist, an NDRI, an SSRI, an S1P modulator, and an AKT activator, across four modalities: snRNA sequencing, bRNA sequencing, proteomics, and electrophysiology, hence prioritizing mechanistic breadth and matched multimodal depth over compound number.^25^ We then used NIS to convert these matched multimodal readouts into category-level and composite drug-response scores and applied the same category framework to external rodent datasets for cross-species comparison. Together, this work establishes NIS as a human-relevant platform for converting multimodal organoid drug-response data into interpretable, category-resolved evidence for CNS drug assessment and cross-species benchmarking.

## Results

### A vascularized human multi-region brain organoid defines the substrate for NIS benchmarking

We first used our previously established MRBO^19^ to define the human tissue context in which NIS would be benchmarked across molecular, cellular, and functional drug-response readouts. We generated MRBOs by differentiating cerebral cortical, endothelial vascular, and brainstem organoids from human iPSCs in parallel, fusing them at day 20, and maintaining the resulting constructs to day 200 (**Figure 1A**). We profiled an extended, longer-aged version of this platform with broader cell-type representation (**Figure 1**) and used this construct as the substrate scored by NIS throughout the study.

**Figure 1.**
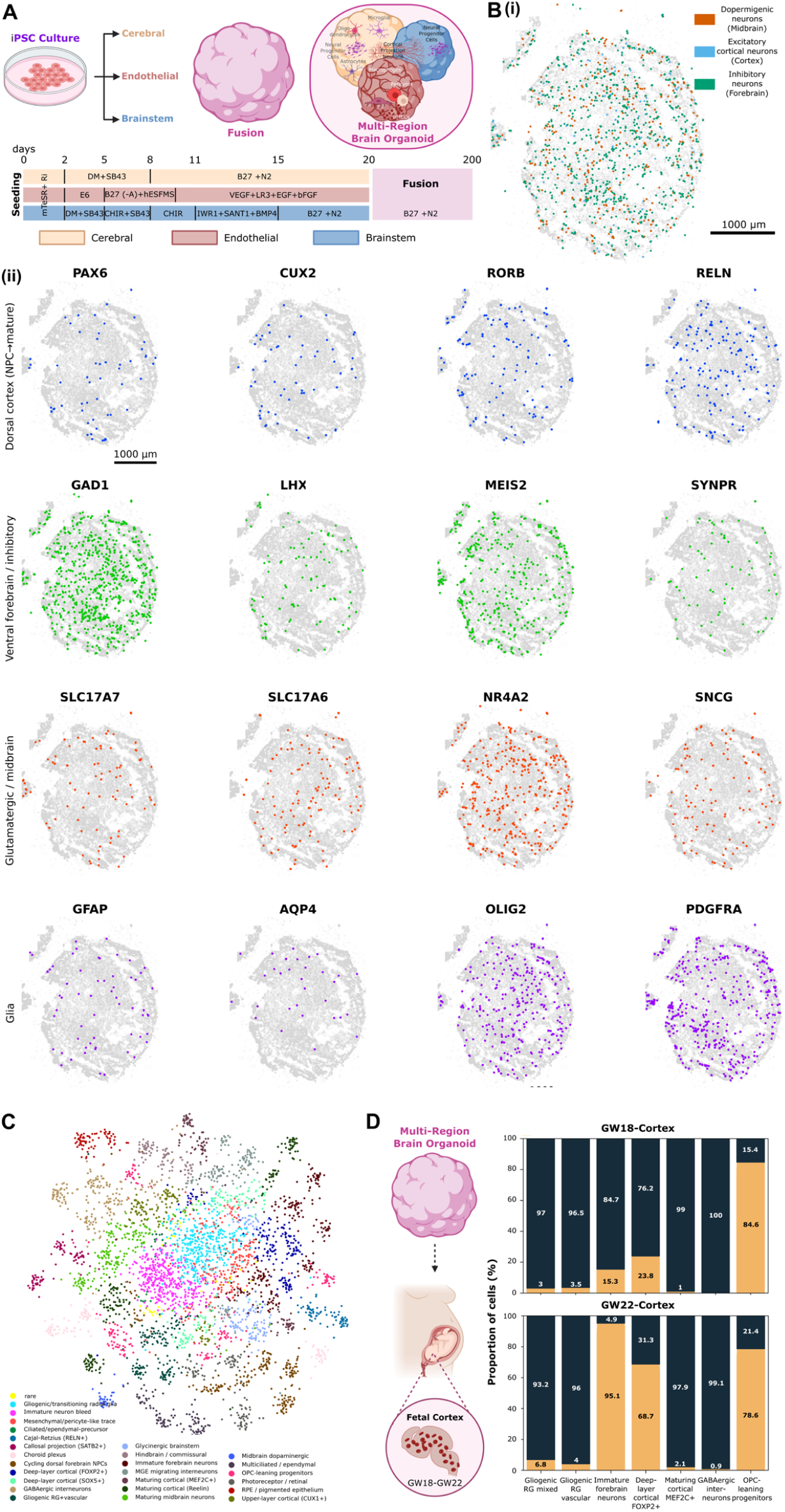
Multi-Region Brain Organoid (MRBO) generation and single-nuclear transcriptomic characterization. (**A**) Schematic of MRBO generation from human iPSCs. Three lineage-specific organoids, cerebral, endothelial, and brainstem, are differentiated in parallel using distinct small-molecule and growth-factor protocols over days 0 to 20, then fused and maintained through day 200 to produce the MRBO. (**B**) Spatial transcriptomic characterization of an MRBO section (**i**) Spatial map of major neuronal classes: midbrain dopaminergic neurons (*orange*), cortical excitatory neurons (*blue*), and forebrain inhibitory neurons (*green*). (**ii**) Spatial expression of 16 canonical markers across four identity groups. Dorsal cortex (*NPC to mature*): PAX6, CUX2, RORB, RELN (n = 116 to 288 spots). Ventral forebrain / inhibitory: GAD1, LHX6, MEIS2, SYNPR (n = 118 to 1,272). Glutamatergic / midbrain: SLC17A7, SLC17A6, NR4A2, SNCG (n = 155 to 557). Glia: GFAP, AQP4, OLIG2, PDGFRA (n = 66 to 808). Colored points are marker-positive cells; *grey* points are all profiled cells. Scale = 1000 µm. (**C**) UMAP of snRNA sequencing nuclei from MRBO colored by annotated cell type (*key, lower left*). 26 annotated cell population span the expected diversity of the construct, including gliogenic / transitioning radial glia, cycling dorsal forebrain NPCs, deep-layer cortical (FOXP2+, SOX5+), upper-layer cortical (CUX1+), callosal projection (SATB2+), Cajal-Retzius (RELN+), maturing cortical (MEF2C+, Reelin), GABAergic and MGE-migrating interneurons, maturing midbrain and midbrain dopaminergic neurons, glycinergic brainstem, hindbrain / commissural, OPC-leaning progenitors, multiciliated / ependymal, mesenchymal / pericyte-like, choroid plexus, photoreceptor / retinal, RPE / pigmented epithelium, and vascular-associated populations. (**D**) Benchmarking of MRBO cell-type proportions against human fetal cortex. MRBO proportions track the GW18 to GW22 ^31^ fetal cortex composition, with dominant mature fractions (84 to 100%) across most categories. This comparison benchmarks the MRBO cortical compartment against age-matched primary tissue only; the midbrain, brainstem, and vascular compartments are not benchmarked here, as no matched primary reference for those regions was included. We pooled replicates from three iPSC lines for spatial/snRNA sequencing sampling (n = 5-6). Panels (**A**) and (**D**) were created in BioRender. Pantula, A. (2026) https://BioRender.com/38co8kh

Using spatial transcriptomic analysis of day-200 sections, we resolved midbrain dopaminergic, cortical excitatory, and forebrain inhibitory neuronal populations, together with dorsal cortical, ventral forebrain inhibitory, glutamatergic/midbrain, and glial marker groups (**Figure 1B (i)**) and sixteen canonical markers separated into dorsal cortical, ventral forebrain inhibitory, glutamatergic/midbrain, and glial groups (**Figure 1B(ii)**).

Using single-nucleus RNA (snRNA) sequencing, we identified 26 annotated cell populations spanning cortical, midbrain, brainstem, glial, choroid plexus, ependymal, retinal, and mesenchymal pericyte-like vascular populations (**Figure 1C**).

To benchmark the developmental state of the cortical compartment, we compared MRBO single-nucleus profiles with mapped human fetal cortex.^31^ We could benchmark only the cortical compartment because matched intact human fetal brain datasets were not available MRBO cortical composition aligned most closely with the gestational week (GW) 18 to GW22 window, with mature fractions of 84 to 100% across matched categories. This placed the extended MRBO in a mid-gestational developmental window suitable for assessing neurodevelopmentally relevant drug responses (**Figure 1D**). Further, by incorporating midbrain and brainstem regions together with an endogenous endothelial compartment, our MRBO broadened the cellular context in which we measured drug responses beyond what cortical-only organoids can capture.

We selected five compounds with dissimilar mechanisms to test whether the MRBO screening platform could resolve drug-specific responses across four multiomics modalities: single-nucleus RNA-seq, bRNA sequencing, proteomics, and electrophysiology. GLP-1 (glucagon-like peptide-1 receptor agonist, GLP-1RA), bupropion (norepinephrine-dopamine reuptake inhibitor, NDRI), fluoxetine (selective serotonin reuptake inhibitor, SSRI), fingolimod (sphingosine-1-phosphate receptor modulator), and SC79 (Akt activator). We then integrated these modality-specific responses with NIS to generate bidirectional biological category scores and a composite drug-response score for each compound.

### Single-nucleus RNA-seq reveals distinct cellular remodeling programs across drug treatments

Out of the four multiomics modalities we first used snRNA sequencing to measure how each compound remodeled the cellular composition and state structure of MRBOs after acute exposure. We treated day-200 MRBOs with GLP-1, Bupropion, Fluoxetine, Fingolimod, SC79, or DMSO for 24 h, then snap-froze the constructs for nuclei isolation and sequencing as described in **Methods, Figure 2**. Although most compounds were selected for direct CNS relevance, we included GLP-1 because it has been reported to have vascular and endothelial activity,^32^ allowing us to test whether the endogenous human endothelial compartment in the MRBO captured endothelial-associated drug responses.

**Figure 2:**
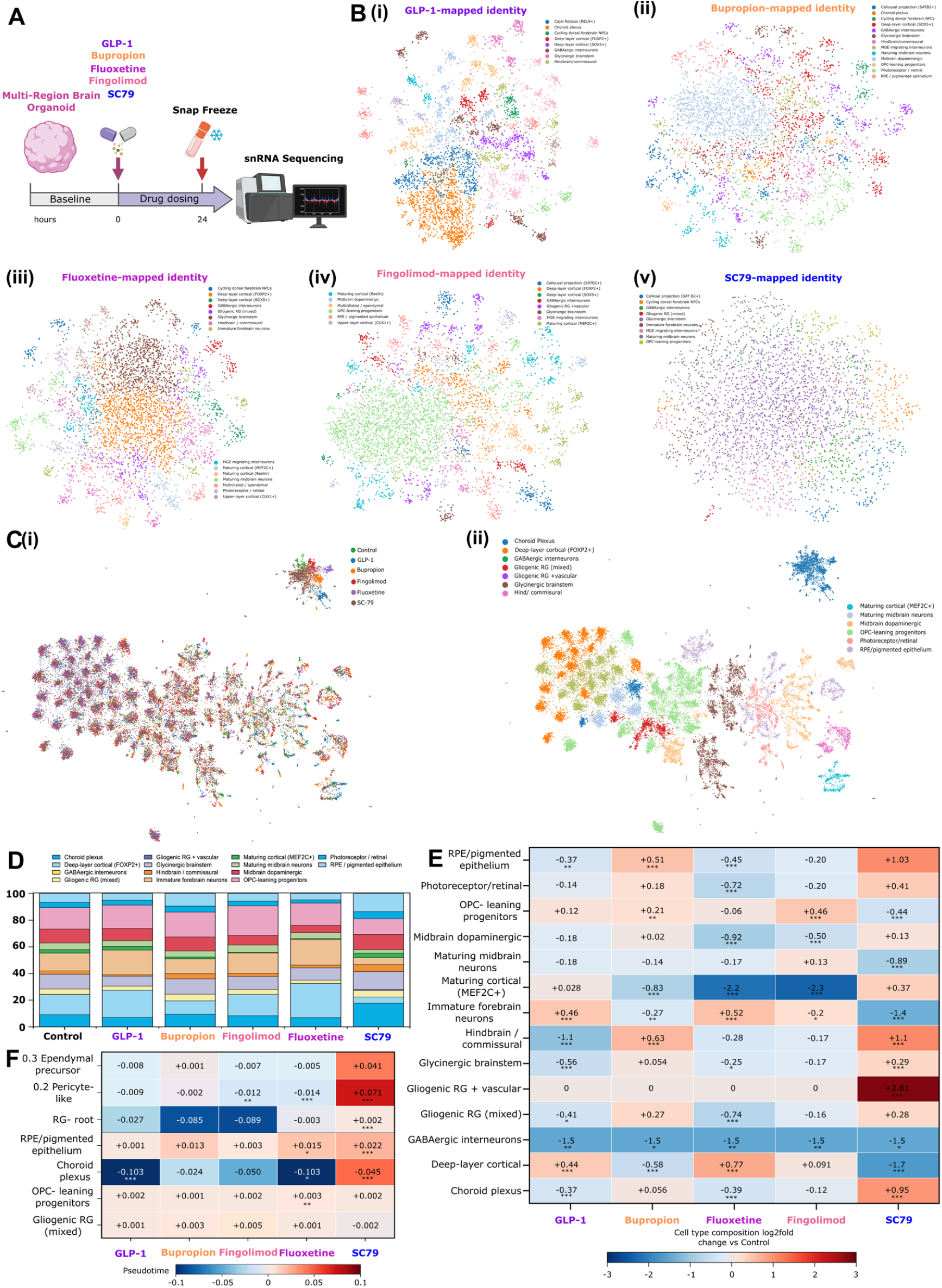
Drug-specific reorganization of MRBOs resolved by snRNA sequencing. (**A**) We treated day-200 MRBOs for 24 h with GLP-1, Bupropion, Fluoxetine, Fingolimod, or SC79, or with vehicle (DMSO; Control). (**B**) We generated per-drug UMAPs colored by mapped identity for all drug treatments. (**C**) Harmony-integrated atlas of all nuclei (n = 24,858) and colored it by drug treatment (*left*) and by the 14 unified cell-type identities used in panels (**D**) to (**F**) (*right*). (**D**) Per-treatment cell-type composition as the percentage of nuclei assigned to each identity. (**E**) Composition shift versus Control as 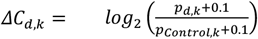 for each drug × identity pair on a diverging *blue-to-red* scale centered at zero. We tested counts by two-sided Fisher’s exact tests on 2 × 2 contingency tables (drug vs Control × identity vs other), with Benjamini-Hochberg (BH) FDR applied across all 66 tests. The full drug × identity table showed strong global association by Pearson’s *χ*^2^ (*χ*^2^ = 2,518.87, degrees of freedom = 65, P < 1 × 10^−300^). (**F**) We computed pseudotime shift versus Control on a shared diffusion-pseudotime (DPT) axis jointly across drug treatments on the Harmony embedding. The DPT root was defined as the Gliogenic-RG cell maximizing the mean scaled expression of SOX2 and NFIA minus the mean scaled expression of the terminal-marker gene set, such that pseudotime increased away from a SOX2^+^, NFIA^+^ radial-glial anchor. Here, *c* denotes a candidate Gliogenic-RG cell, *z*_*c,g*_ denotes the normalized or scaled expression of gene *g* in cell *c*, and *T* denotes the terminal-marker gene set. Each cell *i* was assigned a pseudotime value relative to this root: Each cell shows 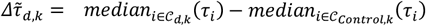 for the indicated identity, on a diverging blue-to-red scale (bound ± 0.103). We partitioned rows into a subcluster trajectory block (Ependymal-precursor, Pericyte-like, RG-root; 15 tests) and a ventricular-lineage block (RPE / pigmented epithelium, Choroid plexus, OPC-leaning progenitors, Gliogenic RG (mixed); 20 tests), for 35 tests total. We compared drug-vs-Control DPT distributions per drug × identity pair by two-sided Mann-Whitney U tests, with BH-FDR applied within each row block. Throughout panels (**E**) and (**F**): ***FDR < 0.001, **FDR < 0.01, *FDR < 0.05; unmarked cells, not significant. For each treatment, we pooled replicates from three iPSC lines for spatial/snRNA sequencing (n = 5-6). Panel (**A**) was created in BioRender. Pantula, A. (2026) https://BioRender.com/38co8kh

We mapped every drug treatment to the annotated MRBO reference and recovered shared, unified cell-type identities across drug backgrounds (**Figure 2B, C**), indicating that the observed compositional changes reflected cell-state redistribution rather than cell-type dropout. Each compound shifted the cellular atlas, defined here as the integrated snRNA sequencing embedding of all nuclei colored by unified cell-type identities, in a distinct pattern **(Figure 2D, E)**. SC79 produced the largest remodeling effect and expanded vascular and progenitor-associated states, including pericyte-like cells, ependymal precursors, RPE/pigmented epithelium, and vascular-signature gliogenic radial glia (global association by Pearson’s *X*^2^: *X*^2^= 2,518.87, degrees of freedom. = 65, P < 1 × 10^−300^; identity-specific shifts tested by two-sided Fisher’s exact tests with Benjamini– Hochberg FDR; **Figure 2E)**. GLP-1 increased immature forebrain and deep-layer cortical FOXP2+ identities, and Bupropion patterned with GLP-1 rather than with the serotonergic or S1P-modulating compounds. Fingolimod and Fluoxetine produced narrower redistributions. At finer resolution, the same split held **(Figure 2G):** GLP-1 and Fluoxetine produced the strongest depletion of choroid plexus identity (both near −0.10; two-sided Fisher’s exact test, BH-FDR < 0.05), whereas SC79 increased RPE/pigmented epithelium (+0.022; two-sided Fisher’s exact test, BH-FDR < 0.05), with each change significant relative to DMSO.SC79 increased RPE/pigmented epithelium (+0.022), with each change significant relative to DMSO.

We anchored a pseudotime trajectory at the radial glia root and placed pericyte-like and ependymal-precursor states downstream of it (**Figure 2F)**. SC79 expanded both downstream states (pericyte-like +0.071, ependymal precursor +0.041) and held the radial glia root nearly steady (+0.002), which indicated a compound that pushed cells outward along the lineage. Bupropion and Fingolimod instead drew the radial glia root down (−0.085 and −0.089) without populating either downstream state, and GLP-1 and Fluoxetine changed it the least. The AKT activator drove cells toward vascular and ependymal fates, whereas Bupropion and Fingolimod depleted the progenitor pool without a corresponding gain in these fates (two-sided Mann - Whitney U tests per drug × identity, BH-FDR < 0.05 within each row block; **Figure 2F)**

### bRNA sequencing and Proteomics define the molecular response streams of the MRBO drug screen

Next, we analyzed the same drug screen using bRNA sequencing and Proteomics from parallel treated organoid samples (**Figure 3**). Details on the methodology and assays used are in the **Methods section**. For the transcriptomic arm, we aligned reads to GRCh38 and identified differentially expressed genes relative to DMSO using DESeq2 (FDR < 0.05, |log2 fold change| > 1). Ranking the top 50 differentially expressed genes by cross-condition variance separated the five compounds into distinct molecular profiles (**Figure 3B**). SC79 produced the most divergent transcriptional profile, with broad upregulation of ribosomal protein genes from the RPL and RPS families and chromatin-associated factors, consistent with AKT-associated translation and proliferative signaling. GLP-1 and Bupropion clustered apart from Fingolimod and Fluoxetine, consistent with their distinct pharmacological targets. Per-drug heatmaps further resolved dose-responsive gene-expression signatures, with many genes scaling from medium to high dose within each compound-specific response (**Figure 3C**).

**Figure 3:**
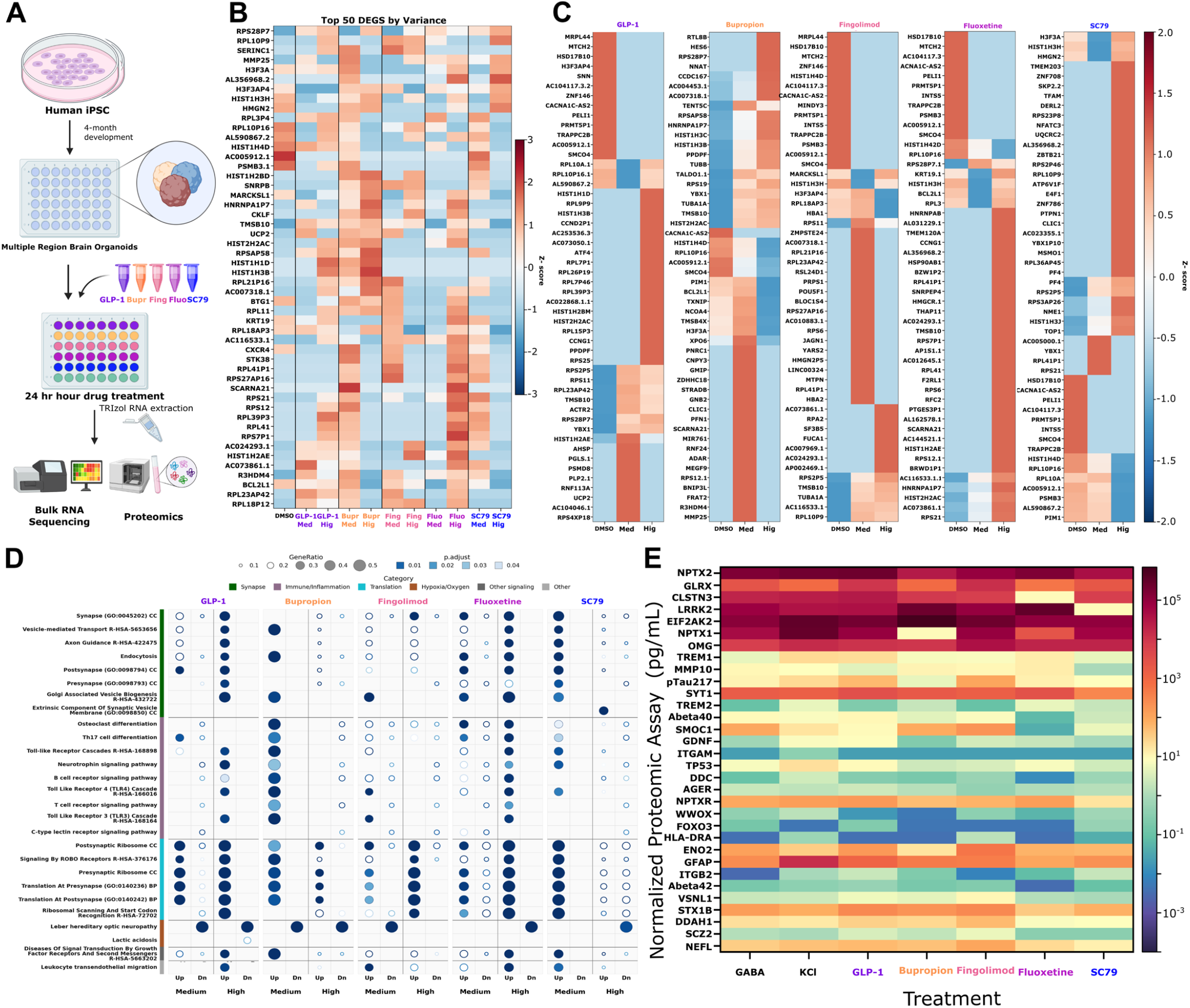
Bulk RNA sequencing and proteomics reveal drug-specific transcriptional and protein-level responses in MRBO. (**A**) We treated MRBOs for 24 hours with five drugs or a DMSO control, then collected samples for both assays. (**B**) Heatmap of the top 50 differentially expressed genes ranked by cross-condition variance of l*o*g_2_(*FPKM* + 1). Rows are individual genes; columns are DMSO and libraries for each drug at each dose (*red*, high; *blue*, low). We ranked 5,189 brain-expressed DEGs observed across all five drugs at both doses by variance across all 11 columns and retained the top 50 genes (variance range, 13.33 - 29.53). We used medium and high doses for bRNA sequencing. **(C)** Per-drug heatmaps of the top 40 brain-expressed DEGs for each compound, ranked by row variance of l*o*g_2_(*FPKM* + 1) across each dose of the drug and DMSO. Per-drug candidate universes: GLP-1, 2,674; Bupropion, 4,775; Fingolimod, 2,189; Fluoxetine, 2,573; SC79, 2,317 genes. Z-scores, color scale (±3), and row clustering are as in (**B**). **(D)** Pathway enrichment dot plot for bulk RNA-seq DEGs from the five drugs at both doses, split into up- and down-regulated gene sets. Enrichment used gseapy Enrichr with Fisher’s exact test and BH adjustment against *GO Biological Process*^33^, *KEGG*^34^, *Human Metabolome*^35^, and *Reactome*^*36*^ restricted to brain-relevant terms (943, 36, and 203 terms, respectively), *SynGO*^37^ and *Allen Brain Atlas*^38^ (up- and down-regulated), *ARCHS4 Tissues*^39^, *GWAS Catalog*^40^, and *DISEASES* ^*41*^. We show the top five terms per drug, dose, and direction after removing anatomy and tissue terms, capped at 35 terms total (27 unique pathways, 273 dots). Dot size denotes GeneRatio, defined as term overlap k/N (range, 0.060 to 1.000; median, 0.241), with open rings for values below 0.25 and filled circles for values at or above 0.25. Dot color denotes adjusted *P* < 0.05 (observed range, 1.6 × 10^−16^ to 5.0 × 10^−2^; median, 2.0 × 10^−3^). (**E**) Proximity extension assay heatmaps. Normalized protein concentrations (pg/mL, l*o*g − *scale* ) for 35 neurology-relevant proteins measured across treatment conditions: GABA, KCl (depolarization controls), and drugs. Proteins include synaptic markers (NPTX2, SYT1, NEFL, STX1B, DDAH1), neuroinflammatory proteins (GFAP, TREM1, TREM2, ITGAM, SMOC1), neurodegeneration-associated proteins (pTau217, Abeta40, BACE1, ENO2, WWOX), and signaling regulators (GLRX, LRRK2, EIF2AK2, FOXO3, ITGB2, AUKM1). We used a high dose for proteomics profiling. For each treatment, we pooled replicates from three iPSC lines for bRNA sequencing (n = 6) and proteomics (n = 6), and from two lines for electrophysiology (n = 3 per condition). To benchmark each assay, we included literature-established positive controls: 100 µM GABA for inhibitory network responses (n = 3) and 30 mM KCl for depolarization-driven cytokine release (n = 3).

We next used pathway enrichment to place the dose-responsive gene sets into biological context (**Figure 3D**). The enriched terms grouped into three broad response classes. The first class captured synaptic and translational programs, including synapse organization, pre- and postsynaptic compartments, synaptic vesicle biogenesis, ROBO receptor signaling, and postsynaptic ribosome and translation-associated pathways. The second class included neurotrophin and BDNF receptor signaling together with potassium channel activity. The third class contained a smaller set of metabolic terms. SC79 again produced the broadest and most significant high-dose enrichment, with strongest representation in translational and synaptic pathways. Using the proteomic panel, we measured 35 neurology-relevant proteins across the five drugs plus GABA and KCl controls (**Figure 3E, SI Figure 12**). We grouped the measured analytes into synaptic proteins (NPTX2, SYT1, STX1B), neuroinflammatory proteins (GFAP, TREM1, TREM2, ITGAM, SMOC1), neurodegeneration-associated proteins (pTau217, Abeta40, BACE1, ENO2, WWOX), and signaling-associated proteins (GLRX, LRRK2, EIF2AK2, FOXO3, ITGB2). SC79 and GLP-1 produced the largest proteomic shifts: SC79 broadly increased synaptic and structural proteins, whereas Fingolimod selectively modulated inflammatory and adhesion-associated proteins. Together, the protein-level responses complemented the bRNA sequencing signatures and showed that drug-associated molecular changes extended beyond transcriptional regulation.

### Network electrophysiology captures functional drug-response signatures of the MRBO

We used MEA recordings as the functional arm of the MRBO drug screen and analyzed extracellular electrical activity as described in Methods. Briefly, we recorded day-200 MRBOs on 48-well multi-electrode arrays, established a 96-h baseline, dosed the constructs at 96, 120, 144, and 168 h, and collected the final recording at 240 h (**Figure 4A**). We focused on spike frequency, network burst duration, spikes per network burst, and burst rate as complementary readouts of spontaneous network activity.

**Figure 4:**
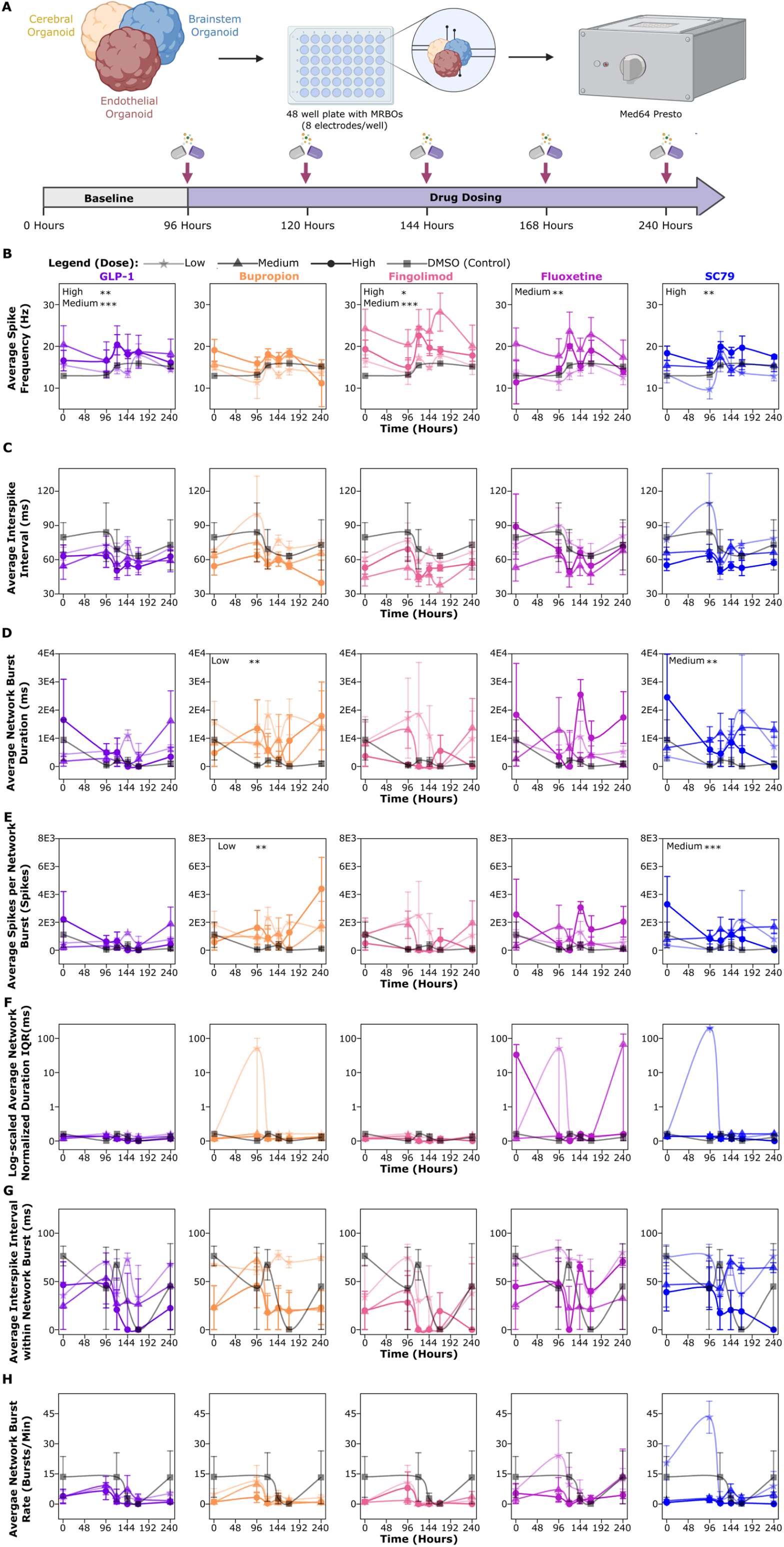
Multielectrode array electrophysiology reveals drug-specific and dose-dependent modulation of MRBO network dynamics over 240 hours. (**A**) Experimental schematic. We plated MRBOs into 48-well MEA plates (8 electrodes per well) and allowed them to establish baseline neural network activity over 96 hours. We initiated drug dosing at 96 hours and continued at 120, 144, and 168 hours, with final recordings at 240 hours across all drug treatments and vehicle control. (**B**) *Average spike frequency (Hz):* GLP-1 (High and Medium), Fingolimod (Medium), Fluoxetine (Medium), and SC79 (High) significantly suppressed spike frequency relative to DMSO, whereas Bupropion did not produce significant changes at any dose (*p < 0.001; **p < 0.01, as indicated). (**C**) *Average interspike interval (ISI, ms)*: Trajectories remained largely stable across dose conditions for most compounds, while SC79 (High) showed the greatest variability, consistent with broader network modulation by AKT activation. (**D**) *Average network burst duration (ms)*. Bupropion (Low) and SC79 (Medium) significantly prolonged network burst duration, with Bupropion showing a progressive increase that peaked between 144 and 192 h (**p < 0.01). (**E**) *Average spikes per network burst*: Bupropion (Low) significantly increased spikes per burst (**p < 0.01), with values rising through 192 h before declining, while SC79 (Medium) also produced a significant increase (***p < 0.001). (**F**) *Log-scaled average network burst duration IQR (normalized)*: Bupropion and Fluoxetine showed transient increases in burst duration variability at intermediate timepoints, whereas GLP-1 and Fingolimod maintained lower variability consistent with network suppression. SC79 (High) had the largest burst-duration IQR, consistent with network destabilization. (**G**) *Average interspike interval within network bursts* (ms). (**H**) *Average network burst rate (bursts/min)*. All data are presented as mean ± SEM. Statistical comparisons are relative to the DMSO control at matched timepoints. For each treatment, we pooled replicates from two lines for electrophysiology (n = 3 per condition). To benchmark each assay, we included literature-established positive controls: 100 µM GABA for inhibitory network responses (n = 3) and 30 mM KCl for depolarization-driven cytokine release (n = 3).

Using one-way ANOVA followed by Dunnett’s multiple comparisons test against DMSO, we found that GLP-1 significantly suppressed spike frequency at high and medium doses (GLP-1 high, p = 0.0068; GLP-1 medium, p = 0.0005), as did Fingolimod at high and medium doses (Fingolimod high, p = 0.0100; Fingolimod medium, p = 0.0007), Fluoxetine at medium dose (p = 0.0031), and SC79 at high dose (p = 0.0016) (**Figure 4B**). Bupropion did not significantly alter spike frequency at any dose. No treatment significantly changed the interspike interval.

Bupropion instead altered burst structure. Low-dose Bupropion significantly increased network burst duration (p = 0.0018) and spikes per network burst (p = 0.0050), consistent with recruitment of larger coordinated bursts rather than faster baseline spiking (**Figure 4D, E**). Medium-dose SC79 produced a similar burst-structure phenotype, increasing network burst duration (p = 0.0045) and spikes per network burst (p = 0.0053) (**Figure 4F**). We did not detect significant treatment effects for normalized network burst duration interquartile range (IQR), interspike interval within network bursts, or network burst rate.

We used these electrophysiological features only for the neurodevelopment axis, as spontaneous network activity reflects circuit maturation and functional organization rather than neuroinflammation, neurodegeneration, or cellular resilience. Thus, MEA provided the functional response stream of the screen, but, like the snRNA sequencing, bRNA sequencing, and proteomic readouts, it described one layer of drug response without independently assigning an overall favorable or unfavorable biological verdict.

### External transcriptomic comparisons reveal gene-level divergence between MRBOs and the rodent brain

Before integrating the multiomic datasets generated from our MRBOs into the four-modality NIS, we tested whether human MRBO drug responses aligned with matched or pathway-related rodent brain transcriptional responses. We tested whether drug-induced transcriptional responses in the human MRBO aligned with matched or pathway-related rodent brain datasets. (**Figure 5A**) This cross-species comparison was important because CNS drug development still relies heavily on animal brain data, whereas human organoids may capture drug-response biology that rodent models do not. At the individual-gene level, MRBO drug responses showed directional divergence rather than broad agreement with rodent brain responses.

**Figure 5:**
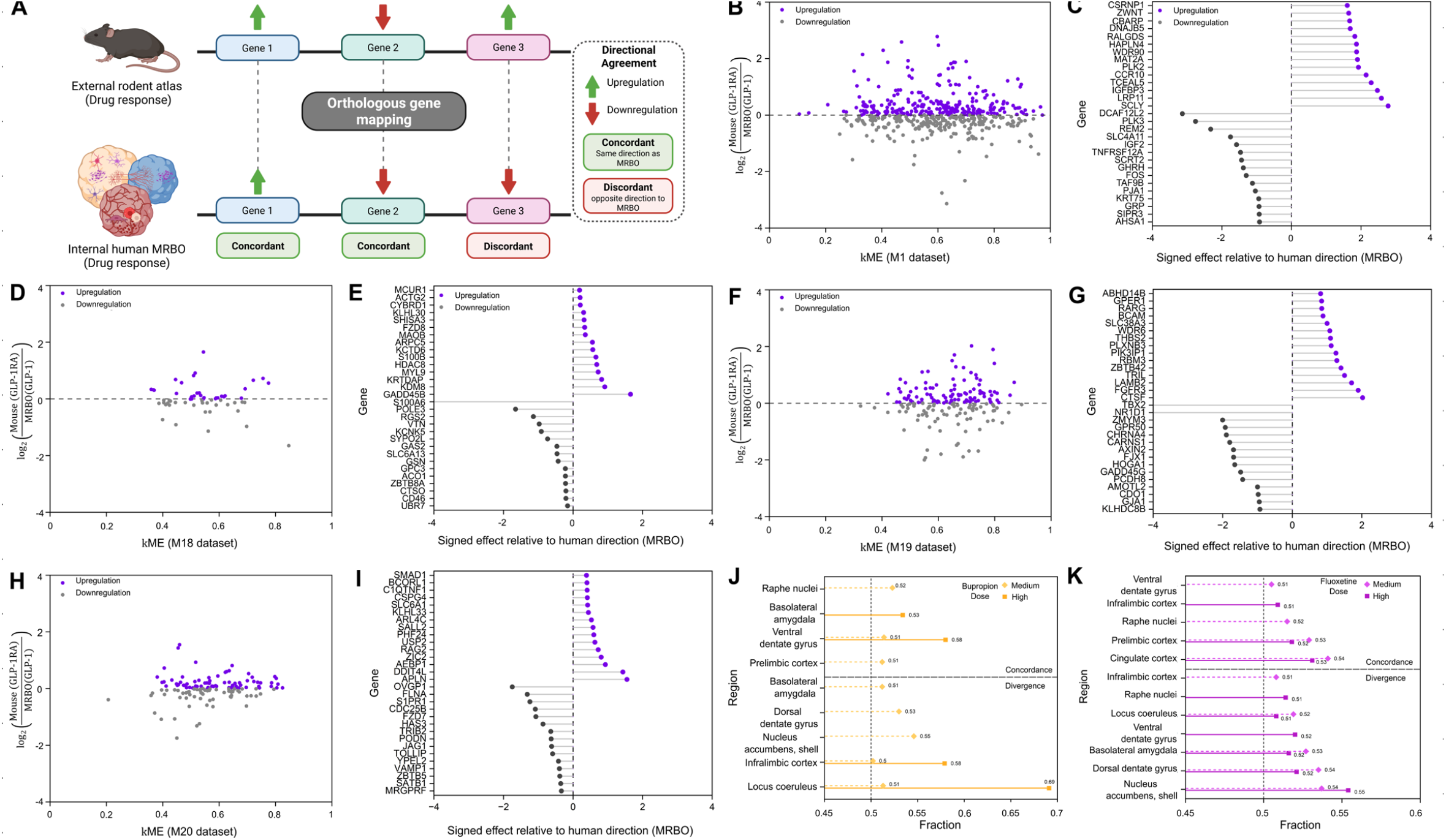
Cross-species concordance of MRBO drug-response signatures with a mouse hindbrain module and a rat brain-region atlas. (**A**) Schematic overview of cross-species directional concordance between rodent atlas and human MRBO drug-response signatures. (**B-I**) Gene-level agreement between GLP-1 effects in MRBO and the M1 co-expression module from a *mouse dorsal vagal complex* snRNA sequencing atlas^42^. We analyzed the GLP-1RA-induced module: *M1* (**B, C**) and the three modules it repressed: *M18* (**D, E**), *M19* (**F, G**), and *M20* (**H, I**). In each scatter, we plotted every module gene detected in the GLP-1 MRBO bulk data as mouse *kME* against the GLP-1 response signed relative to the mouse module direction (positive marked the same direction as the mouse signature, negative the opposite). Each lollipop shows the 15 most concordant and 15 most discordant detected genes per module, which we ranked by signed 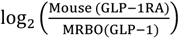 for each gene. (**B**) *M1* dataset (*induced*): GLP-1 MRBO bRNA data,where of 496 genes with defined directional responses in both systems, 256 (51.6%) moved with the mouse signature, which did not differ from chance (binomial p = 0.50; Wilcoxon signed-rank p = 0.40; Spearman ρ = 0.02; Fisher’s exact odds ratio = 0.97). Module centrality did not predict the sign or magnitude of the MRBO response (Spearman ρ = -0.045, P = 0.32; Pearson r = -0.063, P = 0.16). (**C**) The 15 most upregulated and 15 most downregulated *M1* genes detected. (**D**) *M18* dataset (*repressed*): GLP-1 MRBO bRNA data where 28/59 genes moved concordantly, 47.5% (binomial p = 0.79; Wilcoxon p = 0.51; Spearman ρ = -0.04; Fisher’s exact odds ratio = 0.90). (**E**) The 15 most upregulated and 15 most downregulated *M18* genes detected. (**F**) *M19* dataset (*repressed*): GLP-1 MRBO bRNA data, where 97/183, 53.0% (binomial p = 0.46; Wilcoxon p = 0.44; Spearman ρ = 0.06; Fisher’s exact odds ratio = 1.13). (**G**) The 15 most upregulated and 15 most downregulated *M19* genes detected. (**H**) *M20* dataset (*repressed*): GLP-1 MRBO bRNA data, where 68/133, 51.1% (binomial p = 0.86; Wilcoxon p = 0.57; Spearman ρ = 0.14; Fisher’s exact odds ratio = 1.05). (**G**) The 15 most upregulated and 15 most downregulated *M20* genes detected. None of the four modules reached above-chance directional concordance. (**J, K**), We compared the MRBO bulk 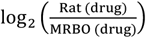 with the atlas documenting the drug response across nine rat brain regions^43^ (GSE194289, bRNA sequencing), for Bupropion (**J**) and Fluoxetine (**K**). For each MRBO drug-by-dose (medium and high) and region pair, we tested the fraction of intersected DEGs with concordant sign against 0.5 (one-sided exact binomial test, BH FDR within each drug and direction). No pair reached FDR-significant concordance for either drug. (**J**), Bupropion diverged significantly at nucleus accumbens shell × Medium (1,274/2,804 = 45.4%, q = 9.4 × 10^−6^), dorsal dentate gyrus × Medium (1,235/2,630 = 47.0%, q = 6.3 × 10^−3^) and locus coeruleus × High (17/55 = 30.9%, q = 1.4 × 10^−2^). (**K**), Fluoxetine diverged significantly at nucleus accumbens shell × High (736/1,650 = 44.6%, q = 1.2 × 10^−4^).

For GLP-1, we compared the MRBO response with GLP-1RA-responsive co-expression modules from the Ludwig dorsal vagal complex (DVC) reference, a GLP-1R-expressing brainstem nucleus and principal site of central GLP-1 agonist action^42^: the single module induced by GLP-1RA (*M1*; **Figure 5B, C**) and the three modules it represses (*M18, M19, M20*; **Figure 5D-I**). For *M1*, among 496 genes with defined directional responses in both systems, 256 moved in the same direction and 240 in opposite directions (51.6% same-direction), a near-symmetric split indicating limited global concordance that was consistent across independent tests (binomial p = 0.50; Wilcoxon signed-rank p = 0.40; Spearman ρ = 0.02; Fisher’s exact odds ratio = 0.97) (**Figure 5B**). The three repressed modules showed the same non-significant pattern: *M18* (28/59 same-direction, 47.5%; binomial p = 0.79; Wilcoxon p = 0.51; Spearman ρ = -0.04; Fisher’s exact odds ratio = 0.90; **Figure 5D**), *M19* (97/183, 53.0%; binomial p = 0.46; Wilcoxon p = 0.44; Spearman ρ = 0.06; Fisher’s exact odds ratio = 1.13; **Figure 5 F**), and *M20* (68/133, 51.1%; binomial p = 0.86; Wilcoxon p = 0.57; Spearman ρ = 0.14; Fisher’s exact odds ratio = 1.05; **Figure 5H**).

We resolved the concordant and discordant genes module by module (**Figure 5C,E,G,I**). Within *M1*, CSRNP1, IGFBP3, LRP11, SCLY, MAT2A, and PLK2 moved in the same direction in both systems, while DCAF12L2, PLK3, REM2, SLC4A11, IGF2, and FOS moved in opposite directions (**Figure 5C**). For M18, MCUR1, ACTG2, CYBRD1, KLHL30, SHISA3, FZD8, MAOB, and GADD45B agreed in direction, whereas S100A6, POLE3, RGS2, VTN, KCNK5, GAS2, SLC6A13, and GSN did not (**Figure 5 E**). M18 had the fewest orthologous genes among the four modules, and the signed effects clustered tightly around zero, except for GADD45B, which showed the largest concordant shift. For M19, ABHD14B, GPER1, RARG, BCAM, SLC38A3, THBS2, PLXNB3, RBM3, LAMB2, FGFR3, and CTSF agreed in direction, whereas TBX2, NR1D1, ZMYM3, GPR50, CHRNA4, AXIN2, GADD45G, PCDH8, GJA1, and KLHDC8B did not, and discordant genes in this module reached larger magnitudes than concordant ones (**Figure 5G**). For M20, SMAD1, BCORL1, C1QTNF1, CSPG4, SLC6A1, SALL2, ZIC2, AEBP1, DDIT4L, and APLN agreed in direction, whereas OVGP1, FLNA, S1PR1, CDC25B, FZD7, HAS3, TRIB2, JAG1, TOLLIP, and SATB1 did not (**Figure 5I**). Developmental patterning genes fell on both sides of the M20 split, so directional agreement did not track gene function within the module.

We compared the bRNA sequencing dataset from our MRBO drug study (Bupropion and Fluoxetine) with an antidepressant rat dataset mapped across nine rat brain regions ^43^ (GSE194289; bRNA sequencing), for Bupropion (**Figure 5J**) and Fluoxetine (**Figure 5K**). Bupropion produced no region that agreed with the rat atlas above chance at either dose (**Figure 5J**). Three regions and dose pairs disagreed. Nucleus accumbens shell at the medium dose gave both the largest intersection and the strongest statistic (1,274/2,804 concordant, 45.4%, q = 9.4 × 10^−6^). Dorsal dentate gyrus followed at the same dose (1,235/2,630, 47.0%, q = 6.3 × 10^−3^). Locus coeruleus fell furthest from chance at the high dose (17/55, 30.9%, q = 1.4 × 10^−2^), though 55 intersected DEGs support that estimate, so we treat its magnitude as provisional and its sign as the informative part. The accumbens and dentate results carry no such caveat. Binomial sampling around a true value of 0.5 would place 45.4% of 2,804 genes roughly five standard errors out, and 47.0% of 2,630 genes roughly three.

Fluoxetine diverged in nucleus accumbens shell at the high dose (44.6%, q = 1.2 × 10^−4^, **Figure 5 K)**. Two explanations account for the lack of directional agreement. Technical factors contributed, including bulk sampling of a whole organoid against regionally dissected rodent tissue, differences in exposure duration, and information lost during one-to-one ortholog mapping. Species of biology contributed as well. Rodent and human neurons differ in receptor subunit composition, in the transcriptional programs that couple activity to gene induction, and in the developmental timing of the circuits these drugs engage, so a compound can produce opposite transcriptional consequences in the two systems without either measurement being wrong. The direction of the divergence favored the second explanation. Purely technical noise would push concordance toward 0.5 and hold it there, yet several region and dose pairs fell significantly below 0.5. Structured anti-correlation of that kind points to real differences in how the two systems respond rather than to measurement error.

### Neural Impact Score (NIS) converts multi-modal MRBO drug responses into bidirectional biological-category scores

While multiomics like snRNA sequencing, bRNA sequencing, proteomics, and electrophysiology, each captured a distinct layer of the cellular, molecular and functional response of the MRBO to the CNS relevant drugs, no individual readout determined whether each compound shifted neural tissue toward circuit formation and maturation, inflammatory or glial stress, degenerative injury, or cellular resilience. We therefore developed the NIS engine to integrate the four multiomics modalities into bidirectional category-level scores for each compound (**Figure 6**).

**Figure 6:**
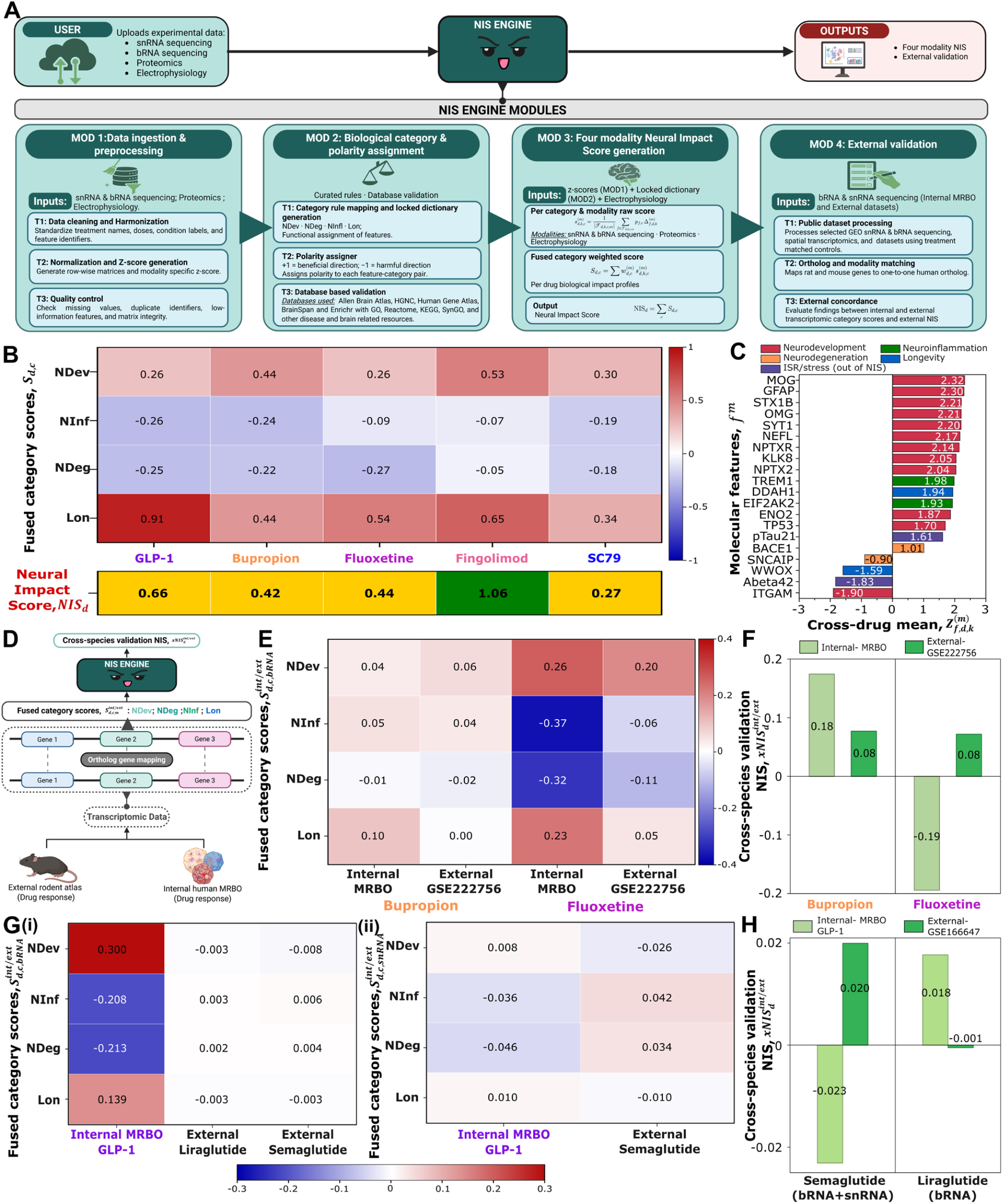
Neural Impact Score engine for multimodal quantification of human-relevant MRBO drug responses and cross-species external validation. **(A)** Overview of the Neural Impact Score (NIS) engine used to indicate the neural impact of a drug across four biological categories (NDev, NDeg, NInf, and Lon). (**B**) Heatmaps displaying fused category scores, *Sd*,_*c*_ for a drug *d*, and biological category *c*, and the neural impact score, *NIS*_*d*_= NIS_*d*_ = *Sd*,_NDev_ + *Sd*,_NInf_ + *Sd*,_NDeg_ + *Sd*,_Lon_. The traffic lights denote the impact *Tier*_*d*_: green, NIS at least 1; yellow, NIS from 0 to< 1; and red, NIS < 0. (**C**) Cross-drug mean standardized responses. (**D**) Schematic of cross-species validation NIS, 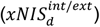, using ortholog-mapped transcriptomic data from internal human MRBO and external rodent atlas drug-response datasets. (**E-F**) NIS-based cross-species transcriptomic validation of antidepressant drug responses across MRBO and rodent brain datasets (**E**) Fused category scores for internal MRBO, 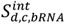 and external GSE222756, 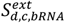 cross-species comparison. We used MRBO bRNA sequencing datasets for Bupropion and Fluoxetine. (**F**) Cross-species validation NIS for MRBO data, 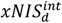 and GSE222756, 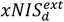. These scores are distinct from the complete four-modality NIS score. (**G-H**) NIS-based class comparison of GLP-1 agonist responses across MRBO and murine dorsal vagal complex datasets. (**G**) Fused category scores for GLP-1 receptor agonist comparisons, reflecting drug-class rather than same-drug validation across datasets. (i) Internal MRBO, 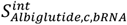 and external liraglutide and semaglutide, 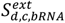 from GSE166647 for the bRNA sequencing datasets. (ii) Internal MRBO, 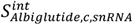 and external liraglutide and semaglutide, 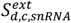 from GSE166647 for the snRNA sequencing datasets. (**H**) Cross-species validation NIS for internal MRBO GLP-1 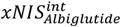 and external Liraglutide and Semaglutide data 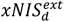 from GSE166647.

We ran four modules (**Figure 6A; Methods**). In Module 1, we ingested electrophysiology, proteomics, bRNA sequencing, and snRNA sequencing datasets, standardized drug names, dose labels, and gene identifiers, and generated modality-specific row-standardized Z-score tables relative to DMSO. In Module 2, we mapped features to four predefined biological categories, Neurodevelopment (NDev), Neuroinflammation (NInf), Neurodegeneration (NDeg), and Longevity (Lon), and assigned each feature a polarity, with positive values indicating favorable movement and negative values indicating unfavorable movement. We locked this dictionary before drug scoring and used database enrichment analyses, including Enrichr^44^, GO^33^, and Reactome^36^, to audit biological coherence rather than tune scores toward expected outcomes (**SI 13)**.

In Module 3, we multiplied each drug-associated z-score difference by its assigned feature polarity, averaged the polarity-adjusted responses within each category and modality, and combined the modality-level scores using fixed, balanced weights. This generated one bidirectional score per biological category and one composite NIS per compound, integrating electrophysiology, proteomics, bRNA sequencing, and snRNA sequencing into a single interpretable framework. We then assigned each composite NIS to a pre-specified tier: GREEN for scores ≥ 1.0, YELLOW for scores from 0 to < 1.0, and RED for scores < 0. In Module 4, we applied the locked category framework to external transcriptomic datasets without modifying the MRBO-derived scores, generating internal and external cross-species validation scores, 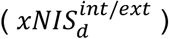, for comparison across compounds and species.

Across the five-drug screen, Fingolimod produced the highest four modality NIS (1.06), driven by positive NDev (0.53) and Lon (0.65), and GLP-1 followed at 0.66 with the highest Lon score (0.91) across the drug panel (**Figure 6C**). Fingolimod cleared the pre-specified GREEN tier. GLP-1, Bupropion (0.42), Fluoxetine (0.44), and SC79 (0.27) fell in the YELLOW tier (**SI Figure 14-17)**.

Feature-level analysis identified the genes and proteins that drove the category scores (**Figure 6C**). MOG, GFAP, STX1B, OMG, SYT1, NEFL, NPTXR, and KLK8 contributed positively within their assigned categories, whereas ITGAM, WWOX, and SNCAIP contributed negatively.

We excluded four integrated stress response proteins, Abeta40, Abeta42, pTau217, and EIF2AK2, from NIS and tracked them as a parallel stress layer rather than scoring them following established integrated stress response frameworks.^45,46^ This design allowed the composite score to capture drug-associated pharmacological signal rather than generic stress burden.

### NIS validates Bupropion and Fluoxetine responses and resolves human-specific GLP-1 pharmacology

We next tested whether NIS could score external datasets without retraining. We distinguished the human-relevant antidepressant responses from our MRBO (Bupropion and Fluoxetine) across a cross-species comparison for rat datasets.^43^ Both compounds showed near-chance directional agreement with rat brain at the individual-gene level (**Figure 5J-K**), their external murine 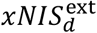 values matched the MRBO scores in sign.

We ran the NIS engine on the dataset obtained from GSE222756^43^ (**Figure 6E, F**) and found that the fused category score, 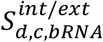 (**Figure 6E**) comparisons showed polarity agreement for NDev for both drugs. While the rest of the categories agreed for Bupropion, NInf, NDeg, and Lon diverged significantly for Fluoxetine, although the polarity remained the same. We also observed that Bupropion scored a positive 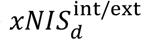 in both the MRBO and external human dataset 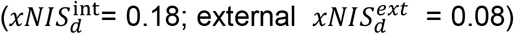. However, Fluoxetine scored positive in the external dataset 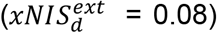 but scored negative in the internal one. 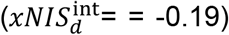 (**Figure 6F**). The latter shows a variance in the drug response across species.

We next applied the same NIS scoring framework to GLP-1 receptor agonist datasets to test whether the framework could support drug-class comparison across human MRBO and murine brain datasets. Because the internal and external datasets contained different GLP-1 receptor agonists, we treated this analysis as drug-class concordance rather than same-drug validation. We used MRBO (GLP-1) and GSE166647 ^42^ (GLP-1R) for Semaglutide and Liraglutide. The Semaglutide dataset comprised two modalities (snRNA and bRNA sequencing) and Liraglutide comprised of one modality (bRNA sequencing).

At the fused category-score level 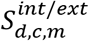, internal GLP-1 showed the strongest bRNA sequencing signal, with positive NDev (0.300) and Lon (0.139) scores but negative NInf (-0.208) and NDeg (-0.213) scores **(SI Figure 17)**. In contrast, the external murine GLP-1 agonist responses showed weaker category-level signal across the same scoring framework, with Liraglutide and Semaglutide producing near-zero bRNA sequencing scores across the available NDeg, NDev, NInf, and Lon categories, ranging from (-0.008) to (0.006) **(SI Figure 16)**. In snRNA sequencing, internal GLP-1 showed smaller category scores, with weak positive NDev (0.008) and Lon (0.010) scores and negative NInf (-0.036) and NDeg (-0.046) scores. External Semaglutide showed a weak, mixed snRNA sequencing profile, with negative NDev (-0.026) and Lon (-0.010), but positive NInf (0.042) and NDeg (0.034).

We then generated the cross-species validation score,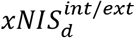. Because Semaglutide included two transcriptomic modalities whereas Liraglutide included only bRNA sequencing, we calculated the internal 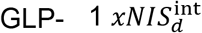 separately for each matched comparison. Internal MRBO GLP-1 produced a negative validation score 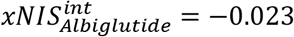 versus the Semaglutide 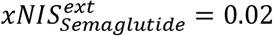. In contrast, it scored positive 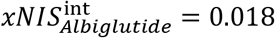 versus the Liraglutide 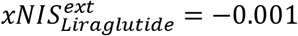.Thus, NIS detected a stronger GLP-1 agonist-associated transcriptomic response in the human MRBO dataset than in the murine dorsal vagal complex comparator datasets.

This divergence likely reflected differences in the cellular compartment captured by each model. GLP-1 receptor agonists act directly on vascular endothelium^32^, and GLP-1 induces transcriptional changes in cultured human endothelial cells. Endothelial-specific deletion of GLP-1R (murine) abolishes vascular effects of human GLP-1 receptor agonism, supporting receptor-dependent endothelial action. As mentioned in the earlier section, because the MRBO contains a substantial human endothelial compartment, we selected GLP-1 a priori to test whether this vascular component would capture endothelial-associated GLP-1 responses that we hypothesized that rodent CNS readouts would miss.

Together, these comparisons showed that NIS could score human MRBO data, independent human transcriptomic data, and rodent brain datasets without retraining or changing the locked feature dictionary. By placing all datasets on the same category-resolved scale, NIS allowed direct comparison of drug responses across compounds, model systems, and species. This was important because the same scoring framework captured positive Bupropion and Fluoxetine responses in human datasets despite weak gene-level concordance with rat brain, while also distinguishing the human MRBO GLP-1 response from near-zero GLP-1 agonist scores in murine dorsal vagal complex datasets. These results support NIS as a quantitative framework for evaluating human-relevant CNS drug responses across experimental paradigms.

## Discussion

We establish the Neural Impact Score as an interpretable scoring framework for human-relevant CNS drug assessment. Using a vascularized day-200 MRBO as the human tissue substrate, we profiled five mechanistically distinct (a GLP-1 receptor agonist, an NDRI, an SSRI, an S1P modulator, and an AKT activator) across four multiomics modalities: snRNA and bRNA sequencing, proteomics, and electrophysiology. CNS-relevant compounds across complex molecular, cellular, and functional readouts and converted these measurements into bidirectional scores within predefined biological categories. In the full MRBO dataset (snRNA & bRNA sequencing, proteomics, and electrophysiology), NIS integrated matched multimodal readouts into category-level and composite scores that separated compound-specific benefits and liabilities. In external rodent datasets, we applied the same locked category definitions in a modality-restricted setting, allowing cross-species comparison even when matched multimodal data were unavailable. Together, these results show that NIS can function both as a multimodal MRBO scoring engine and as a cross-dataset framework for evaluating human-relevant CNS drug responses.

We used our previously established Multi-Region Brain Organoid (MRBO) as the human tissue substrate for this study and allowed it to mature to 200 days. The MRBO provided a vascularized, multi-region human neural model containing cortical, midbrain, brainstem, glial, choroid plexus, ependymal, retinal, and mesenchymal pericyte-like vascular populations. Its cortical compartment aligned most closely with the GW18 to GW22 human fetal cortical window, supporting its use for neurodevelopmentally relevant drug-response assessment. By incorporating midbrain and brainstem regions together with an endogenous endothelial compartment, the MRBO allowed us to evaluate molecular, cellular, vascular, and functional drug responses within the same human substrate.

While multimodal screening usually generates comprehensive molecular, cellular, and functional readouts for each compound, each assay reports only one layer of drug response without giving any interpretation of the holistic impact of the drug on neural tissue. To address this, we developed the Neural Impact Score by converting snRNA/spatial and bRNA transcriptomics, proteomic, and electrophysiological data into bidirectional scores for biological categories like Neurodevelopment, Neuroinflammation, Neurodegeneration, and Longevity. This structure allows us to compare candidate compounds on shared scales in the same biological categories across assay types, model systems, and species.

We used the full MRBO dataset to generate the multimodal NIS. In this setting, NIS integrated all datasets into one bidirectional score per biological category and one composite score per compound. Fingolimod produced the strongest neurodevelopment score (0.53) and remained close to zero or positive across neuroinflammation, neurodegeneration, and longevity respectively, suggesting broad activity across circuit maturation, inflammatory balance, injury-associated signaling, and cellular resilience. This profile may be relevant to neurodevelopmental disorder contexts, including autism spectrum disorder, where circuit maturation and neuroimmune state both contribute to disease biology. GLP-1 showed a different pattern, with the strongest longevity score in the panel (0.91) but a negative neuroinflammation and neurodegeneration score, indicating that GLP-1-linked benefit in this system did not uniformly extend across all categories. Bupropion produced positive neurodevelopment and longevity scores but weaker or negative neuroinflammatory and neurodegenerative profiles, while SC79 and Fluoxetine produced a limited positive neurodevelopment signal but Fluoxetine had a higher longevity score (0.54) with negative neuroinflammation and neurodegeneration scores. These category-specific patterns show why NIS can support rational drug selection beyond composite ranking: drugs with complementary category strengths could be combined to target broader CNS response profiles, while category-specific liabilities could guide exclusion or follow-up testing.

We applied the same category definitions and scoring rules across datasets so that NIS could compare human MRBO and animal brain region drug responses on a shared scale. Because external cross-species datasets rarely contain matched snRNA and bRNA sequencing, proteomics and electrophysiology. We first tested the NIS category framework in a modality-restricted setting. Rather than asking whether the human MRBO reproduced every rodent transcriptional response, we asked whether human organoid and animal brain datasets supported the same directional interpretation within the predefined biological categories. This structure allowed us to evaluate candidate compounds across assay types, model systems, and species, directly addressing the human-relevance problem that drives CNS drug attrition.

The cross-species validation analyses with the antidepressant drugs Bupropion and Fluoxetine directly tested our human-relevance framework. We used the MRBO dataset as the internal human reference, treated the rodent brain datasets as external comparators, and scored each compound across the predefined biological categories using the same locked NIS framework. We then compared the MRBO-derived category scores with the rodent-derived scores (**Figure 6E, F**). At the individual-gene level, both compounds showed near-chance directional agreement between MRBO and rat brain (**Figure 5C, D**). This contrast supports the value of NIS as a category-level comparison framework, because it evaluates whether human and rodent systems support the same biological interpretation without requiring gene-by-gene equivalence.

The GLP-1 cross-species comparison exposed both the advantage of the vascularized MRBO substrate and the limits of rodent neuronal comparators. The two systems inverted sign for both compounds. In the MRBO, Semaglutide scored -0.023 and Liraglutide scored 0.018. In the murine dorsal vagal complex comparator (GSE166647), the ordering reversed: Semaglutide scored 0.020 and Liraglutide scored -0.001. The rodent result is internally coherent, since Semaglutide used two modalities (*bRNA and snRNA*) and the dorsal vagal complex expresses GLP-1R at high density, so the positive external score reflects a genuine neuronal response that maps weakly onto the four defined biological categories in the NIS engine. These results indicate that the MRBO captured different drug-response biology rather than a noisier approximation of the rodent signal. The vascularized MRBO contains an endothelial compartment that neuron-focused rodent comparator datasets cannot capture by design, and this compartment likely contributed to the positive GLP-1 agonist signal observed in the human system. This distinction matters because a change in score direction can alter which compound a screen advances, rather than simply changing the confidence of the ranking. CNS programs involving endothelial signaling, blood-brain-barrier transport, or neurovascular coupling may therefore be mis-prioritized by neuronal comparator datasets, whereas vascularized human organoids provide the relevant cellular context needed to evaluate these mechanisms.

Future studies should extend NIS to patient-derived and disease-context MRBOs, other human-relevant NAM platforms, larger compound panels, and additional CNS-relevant perturbations. Because NIS uses locked category definitions and does not require retraining, researchers can incorporate new datasets on the same scale while preserving comparability across experiments.

Globally, recent regulatory and funding-agency efforts have accelerated the shift toward human-relevant New Approach Methodologies (NAMs),^2,48^ but this shift creates a practical challenge: human-based models must generate evidence that researchers can interpret, compare, and use across experimental systems. This challenge is especially important for CNS drug development, where animal success often fails to predict human outcomes and direct gene-by-gene agreement between human and rodent systems may not be the right benchmark.

This work opens new possibilities for quantitatively interpreting multimodal human-based model systems, where molecular, cellular, and functional readouts are often generated in parallel but interpreted separately. Although we demonstrated NIS using a vascularized MRBO substrate, the framework can extend to other organoids, assembloids, microphysiological systems, tissue chips, and engineered tissue models. We envision NIS as a generalizable scoring framework for human-relevant CNS drug assessment, cross-species benchmarking, organoid-based pharmacology, and next-generation NAM development.

## Materials and Methods

### Multi-region brain organoid generation

We generated Multi-Region Brain Organoids (MRBOs) from established healthy control iPSC lines. The iPSC lines, characterization, passage numbers, and mycoplasma have been previously established.^19^ This study was reviewed by the Johns Hopkins University IRB, which determined that it does not constitute human subjects research under applicable U.S. regulations and thus did not require IRB oversight. Briefly, we differentiated three lineage-specific organoid types: cerebral (cortical), endothelial (vascular), and mid/hindbrain, in parallel and fused them by hanging-drop co-culture on day 20. We then size-matched the embryoid body (EB) diameter at fusion to within ±10% using AggreWell800 plates to ensure reproducible organoid geometry across conditions.

We maintained post-fusion MRBOs in Neurobasal™ medium (Gibco) with added B27+N2 supplements (Gibco) with media changes every 2-3 days and matured them through Day 200 prior to all drug studies. Day 200 MRBOs integrated with a second-trimester human cortex reference atlas^51^, capturing all seven annotated GW22 cell-type clusters and placing the dosing window within the gestational week 16-22 CNS-drug vulnerability window.

### Study Design

#### Drug Dosing and Preparation

We screened five compounds spanning mechanistically distinct CNS pharmacological classes at three dose levels (Low, Medium, High) representing sub-therapeutic, therapeutic, and supra-therapeutic concentrations relative to established plasma *C*_*max*_ values, with concentration brackets accounting for plasma protein binding (84-99%) and brain-to-plasma partitioning (3-10×) following OECD in vitro neurotoxicity convention and previously established pharmacokinetic framework.^52,53^ We have tabulated the doses in **Table 1**.

**Table 1:** Summary of drug dosing scheme used for MRBO treatment experiments.

| Drug / Treatment | Clinical $C_{max}$ | Dose | | |
| --- | --- | --- | --- | --- |
|  |  | Low (L) | Medium (M) | High (H) |
| Bupropion | ≈ 0.5 μM | 0.1 μM | 1 μM | 10 μM |
| Fluoxetine | ≈ 0.5 μM | 0.1 μM | 1 μM | 10 μM |
| Fingolimod | ≈ 10 nM | 1 nM | 10 nM | 100 nM |
| GLP-1 | ≈ 21 nM | 1 nM | 10 nM | 100 nM |
| SC79 | N/A | 1 μM | 5 μM | 10 μM |
| DMSO Vehicle | N/A | 0.1% v/v | 0.1% v/v | 0.1% v/v |

#### MRBO Plating and Experimental Setup

We plated Day 200 MRBOs on 48-well MEA plates (Alpha Med Scientific) and acclimated them for 96 hours with continuous baseline electrophysiological monitoring. We next implemented a repeat-dose regimen to approximate daily dosing. From 96 to 216 hours, we administered the drug every 24 hours after a complete media change. We obtained electrophysiological recordings at five post-dose timepoints (hours 96,120,144,168, 240). For snRNA sequencing, spatial transcriptomics, bulk RNA sequencing, and Proteomics, we treated a separate set of MRBOs at each dose alongside DMSO and collected all molecular samples 24 hours after a single exposure.

### snRNA Sequencing

At 24 h after drug dosing, we transferred MRBOs into Eppendorf tubes after removing most of the residual media. We snap-froze the samples by immersing the tubes in liquid nitrogen in a Dewar for 2 min and immediately stored them at −80°C until RNA-seq processing. Snap-frozen MRBO samples were submitted to AmpSeq, LLC for single-nucleus RNA sequencing.

We processed sn RNA-seq reads using the Illumina DRAGEN Single Cell RNA pipeline with settings appropriate for Fluent BioSciences PIPseq chemistry and snRNA sequencing, including rRNA filtering and intronic-read counting. We aligned and quantified reads against the Illumina hg38-alt_masked rna_v5 reference with GENCODE v44 annotation and exported filtered count matrices in 10x-compatible MTX format. We imported matrices into Scanpy v1.11.5 and concatenated six conditions: DMSO Control, GLP-1, Bupropion, Fingolimod, Fluoxetine, and SC79. We removed nuclei with fewer than 300 detected genes, excluding the top 1% of nuclei by detected gene count as high-count outliers, and discarded genes detected in fewer than five nuclei. After filtering, the dataset contained 24,858 nuclei and 31,843 genes. Because mitochondrial content was negligible across libraries, we did not apply a fixed mitochondrial threshold.

We normalized single-nucleus RNA-seq counts to 10,000 counts per nucleus and applied a log1p transformation. After excluding mitochondrial genes, we identified 2,000 highly variable genes and used this feature set for PCA and Harmony-based dimensionality reduction. We scaled the data with *max_value* = 10 and computed PCA using 50 components. We integrated samples using Harmony to align shared cell-type structures across treatment conditions. Because each sample corresponded to a single treatment condition, sample-level and treatment-level effects could not be separated; therefore, we used Harmony integration only for joint annotation, visualization, and pseudotime analysis. We constructed a neighbor graph from the first 30 Harmony components, followed by UMAP embedding and Leiden clustering.

Leiden clustering at resolution 0.2 produced 31 clusters. We assigned clusters to cell-type identities using curated Control-derived marker panels based on the 22-cluster Control atlas defined in our previously published paper^19^ and Whole Adult Human Brain Census.^54^ For each cluster, we calculated the mean expression for each marker panel and assigned the identity corresponding to the highest mean score. We consolidated these annotations into 14 unified cell-type identities for downstream visualization and analysis. For per-drug and Control-only atlases, we repeated dimensionality reduction, clustering, and marker-panel-based annotation within the relevant subset.

### Pseudotime trajectory, composition, and differential expression analyses

We computed diffusion pseudotime on the Harmony-integrated embedding and analyzed ventricular and gliogenic radial-glia-associated lineage subsets separately. For each subset, we selected a Gliogenic RG root cell using progenitor marker expression and low expression of lineage-specific terminal markers (**SI Figures 1&2**). We compared drug and DMSO Control distributions along the pooled DPT axis using two-sided Mann-Whitney U tests with BH correction (**SI Figures 3&4**).

We assessed treatment-associated shifts in cell-type composition using Fisher’s exact tests comparing each drug and identity against DMSO Control, followed by BH correction. We assessed per-cell-type differential expression between each drug and DMSO Control using Wilcoxon rank-sum tests with BH correction and retained comparisons with at least 10 cells per group (**SI Figure 5-8)**. Because each condition represented a single donor, donor and batch effects could not be separated from treatment effects. We therefore reported composition, expression, and pseudotime analyses descriptively rather than inferential treatment-effect tests.

### Spatial Transcriptomics

We performed targeted spatial transcriptomics on fixed MRBO cryosections using custom padlock and splint probe panels designed against human neural cell-type marker genes. We fixed MRBO samples in 4% paraformaldehyde, cryoprotected them in 30% sucrose, embedded them in OCT, and sectioned them at 25 µm onto silanized and poly-L-lysine-coated coverslips.

We acquired images on a custom fluidics microscopy platform and processed tiled image stacks using illumination correction, 3D deconvolution, fiducial-based registration, pixel-based decoding, stitching, and spot filtering (**SI Figure 9-11**). We analyzed decoded transcript coordinates using *Scanpy* and custom Python scripts. For marker visualization, we plotted curated neural marker genes as spatial scatter plots overlaid on decoded transcripts and generated composite overlays for canonical neuronal markers. Spatial coordinates were reported in full-resolution pixels, and scale bars represent 1000 µm.

### Bulk RNA (bRNA) Sequencing

We extracted total RNA from MRBOs snap-frozen at hour 200 using TRIzol™ reagent (Invitrogen™). Total RNA extraction, sample quality control, poly(A)-enriched strand-specific library preparation, library quality control, and Illumina sequencing were performed by Novogene following the published MRBO protocol.^19^ Novogene returned raw paired-end sequencing data, and downstream analysis was performed in-house.

We trimmed reads for adapter sequences and aligned the clean reads to GRCh38.p14 using STAR. We generated gene-level counts from aligned reads and performed differential expression analysis using DESeq2 with DMSO vehicle as the reference condition. We defined differentially expressed genes as genes with false discovery rate (FDR) < 0.05 and |log2 fold change| > 1.

### Proteomics

We collected conditioned media at 24 hours post-first dose and analyzed it using the Target 48 Neurodegeneration panel (proximity extension assay; 48 nominal assays, Psomagen). We composed proteomic assay wells from 5 MRBOs per condition per iPSC line to ensure adequate analyte concentration. Psomagen performed the assay according to the Olink Target 48 workflow and returned calibrator-normalized protein data for downstream in-house analysis. We mapped 41 proteins to the locked-polarity dictionary (see the *NIS section below*). We reported normalized protein concentrations (to DMSO) on a log NPX scale. All proximity extension assay measurements were performed according to the manufacturer’s protocols.

### Electrophysiology

We used MEA recordings as the functional arm of the MRBO drug screen and analyzed extracellular electrical activity as described in Methods. Briefly, we recorded day-200 MRBOs on 48-well multi-electrode arrays, established a 96 h baseline, dosed the constructs at 96, 120, 144, and 168 h, and collected the final recording at 240 h (**Figure 4A**). We focused on spike frequency, network burst duration, spikes per network burst, and burst rate as complementary readouts of spontaneous network activity.

GLP-1 at medium and high doses significantly suppressed average spike frequency relative to DMSO, as did Fingolimod and Fluoxetine at medium doses, and SC79 at high dose; Bupropion did not significantly alter spike frequency at any dose (**Figure 4B**). Bupropion instead reorganized burst structure, increasing network burst duration at low dose between 144 and 192 h together with a matched increase in spikes per burst, consistent with recruitment of larger coordinated bursts rather than faster baseline spiking (**Figure 4D,E**). SC79 at medium dose similarly prolonged bursts and increased spikes per burst, while high-dose SC79 destabilized network activity (**Figure 4F**). Fluoxetine produced the clearest transient increase in burst rate, peaking between 144 and 168 h (**Figure 4H**).

We used these electrophysiological features only for the neurodevelopment axis because spontaneous network activity reports circuit maturation and functional organization rather than neuroinflammation, neurodegeneration, or cellular resilience. Thus, MEA provided the functional response stream of the screen, but, like the snRNA sequencing, bRNA sequencing, and proteomic readouts, it described one layer of drug response without independently assigning an overall favorable or unfavorable biological verdict.

### Four modality Neural Impact Score Generation

We developed a NIS Engine to integrate snRNA & bRNA sequencing, proteomics, and electrophysiology datasets into a unified drug-response score. We converted each dataset for every drug treatment *d* into a feature-by-drug-treatment matrix, 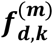 for every modality, *m* for every molecular feature *f* representing genes or proteins and electrophysiology features representing MEA-derived functional metrics. We used the low dose as the primary electrophysiological input, since high-dose exposure sustained across the 240-hour MEA recording would drive toxicity, and the high dose for the 24-hour molecular assays, which captures the maximal acute response before toxicity sets in.

For each modality, we removed features with insufficient numeric values or near-zero variance and applied row-wise z-score normalization across available conditions:

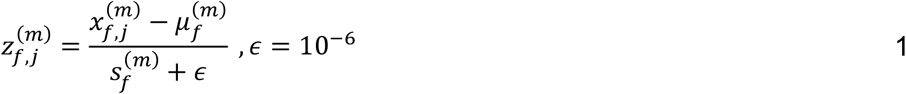

Here, 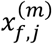 is the processed treatment-level value for feature *f* before standardization, treatment column *j*, and modality *m*. The quantities 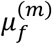 and 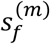 are the mean and sample standard deviation, respectively, calculated across available treatment-level values for that feature. The implementation used the sample standard deviation with Bessel’s correction (*ddof* = 1) and added *ϵ* = 10^−6^ to the denominator for numerical stabilization. Features with fewer than two finite values or a sample standard deviation less than or equal to 10^−6^ were removed before standardization.

We used a curated biological polarity dictionary to assign molecular features to four biological categories (NDev, NDeg, NInf and Lon). Each feature received a polarity value *p*_*f,c*_, where +1 indicated that increased expression relative to control was favorable for that category and −1 indicated expression was unfavorable, thus generating the CNS-relevant locked primary dictionary with category and polarity assignments. Since we used only molecular features for our locked dictionary (snRNA, bRNA sequencing, and proteomics), we assigned the functional features (electrophysiology) separately to the biological category NDev.

For each modality *m*, feature *f*, and drug treatment *d*, and dose *k* we calculated a drug effect relative to control from the normalized values:

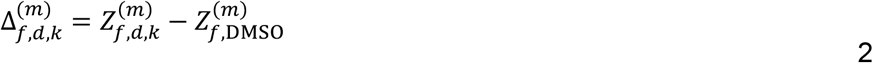

We then calculated per modality raw (**SI Figure 15)** category scores as polarity weighted means of drug-vs-control:

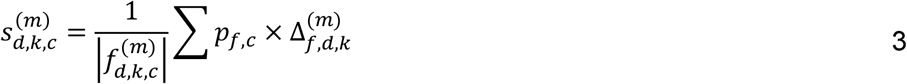

We then calculated the fused category score by assigning each modality a weight 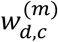 to distribute their impact across biological categories as:

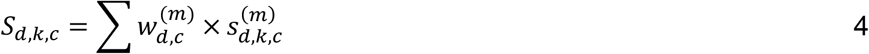

This structure preserved modality separation while generating a single final score per biological category and drug. Finally, we calculated the NIS for each drug by summing the four fused category scores:

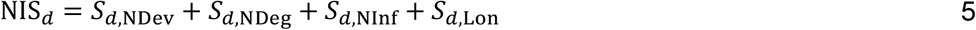

We also assigned our NIS an interpretive color for ease of interpretation

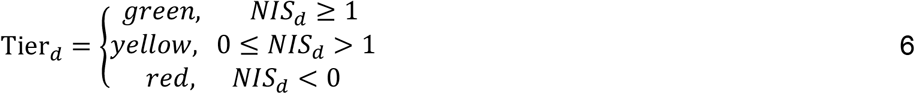

Because we assigned polarity at the feature level, each category score had a consistent direction before summation. Favorable movement increased the category score, and unfavorable movement decreased it. For example, anti-inflammatory movement increased the Neuroinflammation score, whereas inflammatory movement decreased it. We interpreted NIS values as deterministic multimodal favorability scores rather than statistical significance tests.

### External transcriptomic validation

We performed external transcriptomic validation using external datasets (GSE222756, GSE222756, GSE166647) that overlapped with the MRBO drug classes or target pathways. We analyzed each external dataset independently from the internal NIS pipeline and did not use external data to modify the locked MRBO feature dictionary, polarity assignments, modality weights, or final NIS scores. Because the external datasets did not include all four internal modalities, we calculated transcriptomic-only category scores (spatial transcriptomics, snRNA, and bRNA sequencing).

We then *xNIS* values rather than complete four-modality NIS values. We compared external bulk RNA-seq with internal MRBO bRNA sequencing and compared external snRNA sequencing with internal MRBO snRNA sequencing. The spatial transcriptomic analysis was treated as a same-drug, cross-modality comparison with internal MRBO snRNA sequencing. For cross-species datasets, we mapped rat or mouse genes to human dictionary genes using bidirectional one-to-one ortholog mappings and excluded ambiguous or unmapped identifiers.

For external count-based datasets requiring library-size normalization, including GSE222756 and the GSE166648 pseudobulk workflow, we converted raw counts to l*o*g_2_(*CPM* + 1). GSE166647 used the supplied normalized regional bulk RNA-seq values, whereas GSE233571 was processed from Seurat spatial transcriptomic objects. External preprocessing was therefore dataset specific.

For each treatment-control contrast and dataset-specific analysis unit (u), we calculated the external gene-level treatment effect, 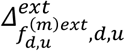 as the mean treatment expression minus the corresponding mean control expression. We used the 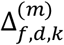 from module 3 for the internal MRBO dataset. We then calculated each external and internal (MRBO) biological-category score, 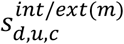, applied modality weights of 0.5 each (in case of Semaglutide data), and calculated the fused category score 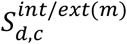.

We calculated the external transcriptomic validation score 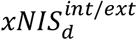 by aggregating the valid external category scores:

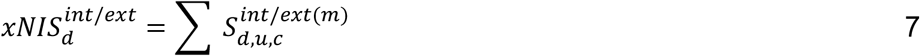

We also assessed directional concordance between MRBO and external datasets at the gene and category levels using binomial tests, Wilcoxon signed-rank tests, one-sample t-tests, Mann-Whitney U tests, Fisher’s exact tests, and Pearson or Spearman correlation, as appropriate for each comparison. We treated external transcriptomic scores as validation readouts of directional concordance rather than as replacements for the full four-modality NIS for the drug-treated MRBOs.

### Rigor and reproducibility

We ran DMSO vehicle controls on every plate across all four modalities. For each treatment, we pooled replicates from three iPSC lines for spatial/snRNA sequencing (n = 5-6), bRNA sequencing (n = 6), and proteomics (n = 6), and from two lines for electrophysiology (n = 3 per condition). To benchmark each assay, we included literature-established positive controls^19^: 100 µM GABA for inhibitory network responses (n = 3) and 30 mM KCl for depolarization-driven cytokine release (n = 3).

All statistical tests were two-sided unless otherwise specified. We performed appropriate parametric or non-parametric tests as specified by the analysis panel for the electrophysiology data; comparisons to the DMSO vehicle at matched timepoints were performed. For snRNA sequencing composition analyses, we used two-sided Fisher’s exact tests with BH correction across 66 tests. For bRNA sequencing differential expression, DESeq2 was applied with FDR < 0.05 and |log2 fold change| > 1 as the significance threshold. For cross-species directional concordance analyses, the statistical frameworks employed included two-sided binomial tests, Wilcoxon signed-rank tests, one-sample t-tests, Mann-Whitney U tests, Spearman and Pearson correlation analyses, and Fisher’s exact test, as described in the External Cross-Species Validation section above. NIS tier thresholds were pre-specified prior to data analysis and committed to version-controlled repositories before unblinding; they were not adjusted post hoc. All analyses were performed in Python and R. Significance thresholds were defined as *p < 0.05, **p < 0.01, and ***p< 0.001.

## Author Contributions

**AP**: Conceptualization, data curation, formal analysis, investigation, methodology, validation, visualization, writing – original draft, writing – review & editing **VS, OS and NL**: Data curation, formal analysis, investigation, methodology, validation, visualization, writing – original draft, writing – review & editing; **KJ**: Methodology, writing – review & editing; **RP&JS**: Investigation, methodology, validation, visualization, data curation for Spatial Transcriptomics, writing – review & editing; **GSO’B**: Supervision, investigation, methodology, validation, visualization, data curation for Spatial Transcriptomics, writing – review & editing; **AK**: Funding acquisition, supervision, conceptualization, formal analysis, methodology, validation, visualization, writing – original draft, writing – review & editing.

## Ethics declarations

A.K. is a co-founder and equity holder of Organotics, Inc., which has licensed intellectual property related to the technology described in this work from Johns Hopkins University. A.P, O.S and A.K are inventors on a patent application related to this technology filed by Johns Hopkins University (Provisional Patent Application No. 64/135,767). The remaining authors declare no competing interests.

